# Genetic and pharmacological manipulations of glyoxalase 1 mediate ethanol withdrawal seizure susceptibility in mice

**DOI:** 10.1101/2020.11.19.389452

**Authors:** Amanda M. Barkley-Levenson, Amy Lee, Abraham A. Palmer

**Affiliations:** Department of Psychiatry, University of California San Diego, La Jolla, CA 92093, USA; Institute for Genomic Medicine, University of California San Diego, La Jolla, CA 92093, USA

**Keywords:** ethanol, withdrawal, genetics, handling induced convulsions, glyoxalase 1, GABA

## Abstract

Central nervous system (CNS) hyperexcitability is a clinically significant feature of acute ethanol withdrawal. There is evidence for a genetic contribution to withdrawal severity, but specific genetic risk factors have not be identified. The gene glyoxalase 1 *(Glo1)* has been previously implicated in ethanol consumption in mice, and GLO1 inhibition can attenuate drinking in mice and rats. Here, we investigated whether genetic and pharmacological manipulations of GLO1 activity can also mediate ethanol withdrawal seizure severity in mice. Mice from two transgenic lines overexpressing *Glo1* on different genetic backgrounds (C57BL/6J [B6] and FVB/NJ [FVB]) were tested for handling induced convulsions (HICs) as a measure of acute ethanol withdrawal. Following an injection of 4 g/kg alcohol, both B6 and FVB mice overexpressing *Glo1* showed increased HICs compared to wild type littermates, though only the FVB line showed a statistically significant difference. We also administered daily ethanol injections (2 g/kg + 9 mg/kg 4-methylpyrazole) to wild type B6 mice for 10 days and tested them for HICs on the 10^th^ day, following treatment with either vehicle or a GLO1 inhibitor (S-bromobenzylglutathione cyclopentyl diester [pBBG]). Treatment with pBBG reduced HICs, although this effect was only statistically significant following two 10-day cycles of ethanol exposure and withdrawal. These results provide converging genetic and pharmacological evidence that GLO1 can mediate ethanol withdrawal seizure susceptibility. We discuss the possible role of GLO1’s substrate, methylglyoxal, acting as an agonist at GABA_A_ receptors.

## 1. Introduction

Alcohol (ethanol) withdrawal is a key feature of alcohol dependence and includes physiological and affective symptoms such as tremors, nausea, vomiting, irritability, insomnia, and anxiety. Withdrawal symptoms are included in the diagnostic criteria for an alcohol use disorder [1], and withdrawal may be related to relapse risk [e.g. 2–4]. In severe cases, acute withdrawal can lead to serious consequences such as delirium, seizures, or even death [5–7]. The determinants of ethanol withdrawal severity are not fully known, but there are likely multiple contributing factors including drinking history, past withdrawal experience, co-occurring use of other drugs, structural brain lesions, and genetics [5, 8]. Understanding the biological factors that confer risk for severe ethanol withdrawal may therefore lead to improved prevention and treatment options.

There is considerable evidence from both human and model organism studies for genetic contributions to ethanol withdrawal severity [9–13]. In humans, gene polymorphisms associated with many neurotransmitter systems have been explored in relation to ethanol withdrawal (for review see [8]). Several of these candidate gene and association studies have identified relationships between specific genetic variants and ethanol withdrawal severity [14–18]. However, large-scale human genetics studies have predominately focused on alcohol consumption and alcohol use disorder diagnosis, rather than withdrawal severity. Animal models are a useful complementary tool for studying novel genetic contributions to ethanol related traits, since animal models can isolate specific aspects of ethanol withdrawal such as seizure susceptibility. Heritability of susceptibility to ethanol withdrawal seizures has been well-established in mouse models: inbred mouse strains and recombinant inbred panels show a high degree of phenotypic variation for this trait [11, 19–21], and selective breeding has produced withdrawal seizure prone and resistant lines of mice [10, 22].

Here, we use a mouse model of ethanol withdrawal to investigate the role of glyoxalase 1 (GLO1) in withdrawal seizure severity. GLO1 is a ubiquitously expressed enzyme that metabolizes the glycolytic byproduct methylglyoxal (MG). We have shown previously that MG can act as a partial agonist at GABA_A_ receptors and that inhibition of GLO1 leads to increased levels of MG [23]. Genetic and pharmacological manipulations of the GLO1 system, as well as direct administration of MG, have been associated with changes in anxiety- and depression-like behaviors, ethanol consumption, and seizure threshold. Specifically, decreased anxiety-like behavior [23, 24], decreased depression-like behavior [25, 26], decreased ethanol consumption [27, 28], and decreased susceptibility to pilocarpine- and picrotoxin-induce seizures [29] have all been seen following GLO1 inhibitor treatment. Some of these effects are also produced by genetic knockdown of *Glo1* or direct MG administration. In contrast, *Glo1* overexpression produces opposite effects (i.e. increased anxiety-like behavior and ethanol consumption) [23, 25, 27]. Furthermore, *Glo1* overexpressing mice show increased GLO1 enzymatic activity and decreased whole brain MG levels, whereas wild type mice treated with a GLO1 inhibitor show the opposite effects [23].

Significant changes in GABAergic signaling following chronic ethanol use are believed to underlie in part the central nervous system hyperexcitability seen in ethanol withdrawal (e.g. [30–32]). We hypothesize that modulating GLO1 activity could impact ethanol withdrawal, potentially through downstream effects on MG levels and subsequent GABAergic activation. In the present studies, we examined whether overexpression of *Glo1* could potentiate ethanol withdrawal seizure severity, and whether pharmacological manipulation of GLO1 activity by treatment with a GLO1 inhibitor could attenuate ethanol withdrawal seizure severity in mice.

## 2. Materials and Methods

### Animals and husbandry

All *Glo1* transgenic mice used in Experiments 1 and 2 were bred in house. Generation of the *Glo1* transgenic mice by insertion of a BAC transgene has been previously described [23]. The *Glo1* transgenic lines are maintained by breeding male mice heterozygous for the *Glo1* overexpression to wild type female mice (B6 or FVB depending on the line). Resulting heterozygous and wild type littermates were used as experimental animals. For Experiment 3, wild type B6 mice were purchased from the Jackson Laboratory (Bar Harbor, ME). For all experiments, mice were housed 2-5 per cage on wood chip bedding and food (Envigo 8604, Indianapolis, IN) and water were provided *ad libitum.* Mice were maintained on a 12hr/12hr light/dark cycle with lights on at 06:00. Behavioral testing was conducted during the light phase. All procedures were approved by the University of California San Diego Institutional Animal Care and Use Committee (#S15226) and were conducted in accordance with the NIH Guidelines for the Care and Use of Laboratory Animals.

### Handling induced convulsions

induced convulsions (HICs) are used to assess susceptibility to ethanol withdrawal seizures and serve as a measure of CNS hyperexcitability. We used the 7-point scale developed by Crabbe and colleagues (e.g. [33]) based on the original procedure by Goldstein and Pal [34]. Briefly, mice are lifted by the tail and observed for convulsion signs. If lifting alone does not produce a convulsion, the mouse is gently spun in a 360-degree arc. Scores range from 0 (no convulsion) to 7 (severe and sometimes lethal seizure due to environmental stimuli). Baseline HIC assessments are made prior to ethanol administration. In Experiments 1 and 3, HICs were assessed hourly from 2-10 hours after the final ethanol injection. In Experiment 2, HICs were assessed only during peak withdrawal (6-8 hr after the final ethanol injection). In all experiments, a final HIC measurement was taken at 24 hr into withdrawal. Experimenters were blinded to genotype and treatment for all HIC assessments, and the same experimenter performed all HIC assessments within a given experiment. A withdrawal severity score was calculated by summing the total post-ethanol HIC scores to find the area under the curve (AUC) of the HIC time course in Experiments 1 and 2, and the HIC scores from hours 4-10 in Experiment 3 [35, 36]. Peak HIC score was determined by averaging the highest HIC score recorded for each animal with the HIC scores from the hourly assessments immediately preceding and following the time of maximum score [21]. In instances where there were multiple time points with the same maximum score, the earliest occurrence was used. Latency to peak HIC was the time at which the highest score first occurred. Animals that did not show any HICs were excluded from the latency analyses.

### Drugs

The GLO1 inhibitor S-bromobenzylglutathione cyclopentyl diester (pBBG) was synthesized in the laboratory of Alexander Arnold at the University of Wisconsin Milwaukee, as previously described [27]. The dose of 25 mg/kg was chosen because it was believed to be an intermediate dose in the behaviorally-active range; 12.5 mg/kg and 50 mg/kg doses have been shown to reduce ethanol drinking and anxiety-like behavior, respectively in previous studies [23, 27], and 25 mg/kg can attenuate withdrawal-associated escalation of ethanol drinking in dependent rats [28]. pBBG was dissolved in vehicle (8% DMSO/18% tween80/saline) and administered i.p. (injection volume 0.01 ml/g). Ethanol (Deacon Laboratories Inc., King of Prussia, PA) was mixed in saline (20% v/v) and administered i.p. at a dose of 4 g/kg in Experiments 1 and 2. In Experiment 3, the alcohol dehydrogenase inhibitor 4-methylpyrazole (9 mg/kg; Sigma-Aldrich, St. Louis, MO) was dissolved in 20% ethanol solution or physiological saline, such that animals received only a single injection for both the ethanol or saline and pyrazole administration.

### Experiment 1: Acute ethanol withdrawal in Glo1 transgenic mice on a B6 background

Male and female mice from the B6 background *Glo1* transgenic line were used in this experiment (n=4/sex/genotype). All animals were between 59-153 days at the start of testing. Mice were weighed, assessed for baseline HIC score, and then injected i.p. with 4 g/kg ethanol and returned to the home cage. Starting two hours after the ethanol injection, mice were assessed hourly for HICs with the final HIC assessment made at 10 hr after injection. The following morning, 24 hr after the previous injection, the final HIC assessment was made.

### Experiment 2: Acute ethanol withdrawal in Glo1 transgenic mice on an FVB background

Because B6 background mice in Experiment 1 showed limited ethanol withdrawal seizure susceptibility, we sought to extend our studies to include FVB *Glo1* transgenic mice because FVB mice have higher HIC scores during ethanol withdrawal than B6 [21]. Peak withdrawal scores were seen from 6-8 hr after injection in Experiment 1, so in Experiment 2 we limited HIC assessments to this peak withdrawal window. Only females were used for Experiment 2 due to animal availability (n=9-12/genotype); all mice were between 241-287 days of age at testing. As in Experiment 1, mice were weighed, given a baseline HIC assessment, and then injected with 4 g/kg ethanol and returned to their home cages. HICs were assessed at 6, 7, and 8 hr after injection and again the next morning at 24 hr after injection.

### Experiment 3: GLO1 inhibitor effects on chronic ethanol withdrawal

To assess whether pharmacological manipulations of GLO1 activity could also influence ethanol withdrawal, we tested whether the GLO1 inhibitor pBBG could attenuate withdrawal seizure severity following chronic ethanol exposure. Importantly, GLO1 inhibition should produce effects that are opposite to those produced by *Glo1* overexpression. Male B6 mice were used for this experiment (n=11-13/treatment group); all animals were 75-77 days of age at the start of testing. Male B6 animals were chosen because the majority of our studies examining GLO1 inhibitor effects on behavior and effective dose ranges have used male B6 mice. Because wild type B6 animals showed limited withdrawal seizure susceptibility following acute ethanol injection in Experiment 1, we used a chronic ethanol dosing model in this experiment to induce dependence. Specifically, we used a 10-day ethanol injection paradigm adapted from Perez and De Biasi [37] that has been shown previously to produce somatic and affective signs of withdrawal in male B6 mice. Mice received a daily injection of either 2 g/kg ethanol (withdrawal group) or 0.9% saline (non-withdrawal control group) with 9 mg/kg 4-methylpyrazole for 10 days.

Baseline HICs were assessed before the first injection was given. On Day 10, HIC time course was assessed starting 2 hr after the final injection and again hourly until 10 hr post-injection. At 3.5 hr after the final ethanol injection, mice received an i.p. injection of either 25 mg/kg pBBG or vehicle. This pretreatment time was chosen to give sufficient time for GLO1 inhibition to lead to increased MG levels during peak withdrawal. A final HIC assessment was made the next day at 24 hr after the final ethanol injection.

To determine whether the effectiveness of pBBG changed with increased ethanol exposure, and due to low levels of withdrawal seizure susceptibility seen following the initial ethanol exposure, we repeated the same procedure for a second cycle of ethanol exposure and withdrawal. Following the first 10-day ethanol withdrawal cycle, mice were allowed to remain undisturbed for 6 weeks in their home cages before we repeated the 10-day procedure described above. All mice received the same drug treatment and the same experimental procedures were used in both ethanol withdrawal cycles, including a re-baselining HIC assessment before the start of the second ethanol exposure period.

### Statistical analyses

Data are expressed as mean ± SEM. All statistical analyses were performed using SPSS (version 27; IBM Corp, Armonk, NY). HIC time course data were analyzed by mixed-model analysis of variance (ANOVA) with a repeated measure of time point and between-subjects factors of genotype, treatment group, and/or sex, as relevant for the given experiment. A Huynh-Feldt correction for violations of sphericity was used for repeated measures analyses. HIC AUC, peak HIC score, and latency to peak HIC score data were analyzed by one- or two-way ANOVA. Significance was set at α=0.05 for all tests. Individual-level data can be found in the supplement in Table S1 (total number of mice per maximum HIC score in each experiment).

## 3. Results

### Experiment 1: Acute ethanol withdrawal in Glo1 transgenic mice on a B6 background

Withdrawal HIC time course (**Fig. 1a**) was analyzed using a mixed-model ANOVA with a repeated measure of time and between-subjects factors of genotype and sex. There were statistical trends toward main effects of time (*F*_5.83, 70_=1.95, p=0.086) and genotype (*F*_1,12_=4.2, p=0.063), and a main effect of sex (*F*_1,14_=4.2, p=0.022, males>females). There were no significant interactions between any factors (p>0.1 for all). HIC time course, AUC, and peak HIC scores for males and females (collapsed on genotype) are shown in supplemental figure S1. Data in Figure 1 are shown broken down by genotype to allow for comparison with the results of Experiment 2.

**Figure 1.**
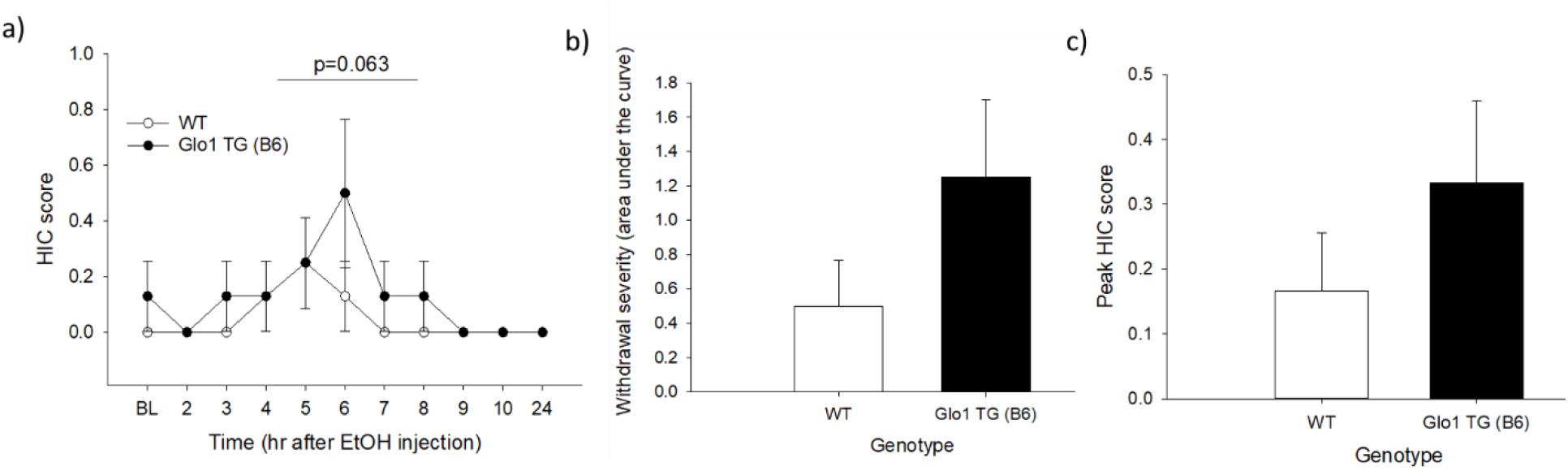
Acute ethanol withdrawal as assessed by handling-induced convulsions (HICs) in wild type (WT) and *Glo1* overexpressing (TG) mice on a B6 genetic background. Panel **a** shows time course of withdrawal, with baseline HIC measurements made immediately before ethanol injection. There was a statistical trend toward a main effect of genotype, with *Glo1* overexpressing mice trending toward higher HIC scores across the time course. Panel **b** shows cumulative HIC scores (area under the curve, AUC) for each genotype. Panel **c** shows peak HIC score for each genotype. Genotype differences for both AUC and peak HIC score were not statistically significant. N=8/genotype.

To determine differences in overall withdrawal severity, we analyzed area under the HIC curve. HIC AUC did not differ significantly between the genotypes (*F*_1,12_=2.35, p=0.151; **Fig. 1b**), though there was a statistical trend towards a main effect of sex (*F*_1,12_=4.17, p=0.064; **Fig. S1b**). There were no main effects of either genotype (*F*_1,12_=1.26, p=0.283; **Fig. 1c**) or sex (*F*_1,12_=2.84, p=0.118; **Fig. S1c**) for peak HIC score (**Fig. 1c**). Latency to peak HIC also showed no main effects of sex or genotype (*F*_1,5_≤0.091, p≥0.771, data not shown). There were no significant genotype x sex interactions for any measure (p>0.1 for all).

### Experiment 2: Acute ethanol withdrawal in Glo1 transgenic mice on an FVB background

Withdrawal HIC time course (**Fig. 2a**) was analyzed using a mixed-model ANOVA with a repeated measure of time and a between-subjects factor of genotype. There were main effects of time (*F*_3.88,73.68_=37.02, p<0.001) and genotype (*F*_1,19_=6.32, p=0.021), but no significant time x genotype interaction. Analysis of HIC AUC (**Fig. 2b**) and peak HIC score (**Fig. 2c**) showed that transgenic mice had significantly greater withdrawal severity (*F*_1,19_=5.24, p=0.034) and higher peak HIC scores (*F*_1,19_=4.99, p=0.038) than wild type mice. Latency to peak HIC did not differ between the genotypes (*F*_1,18_=0.015, p=0.903, data not shown).

**Figure 2.**
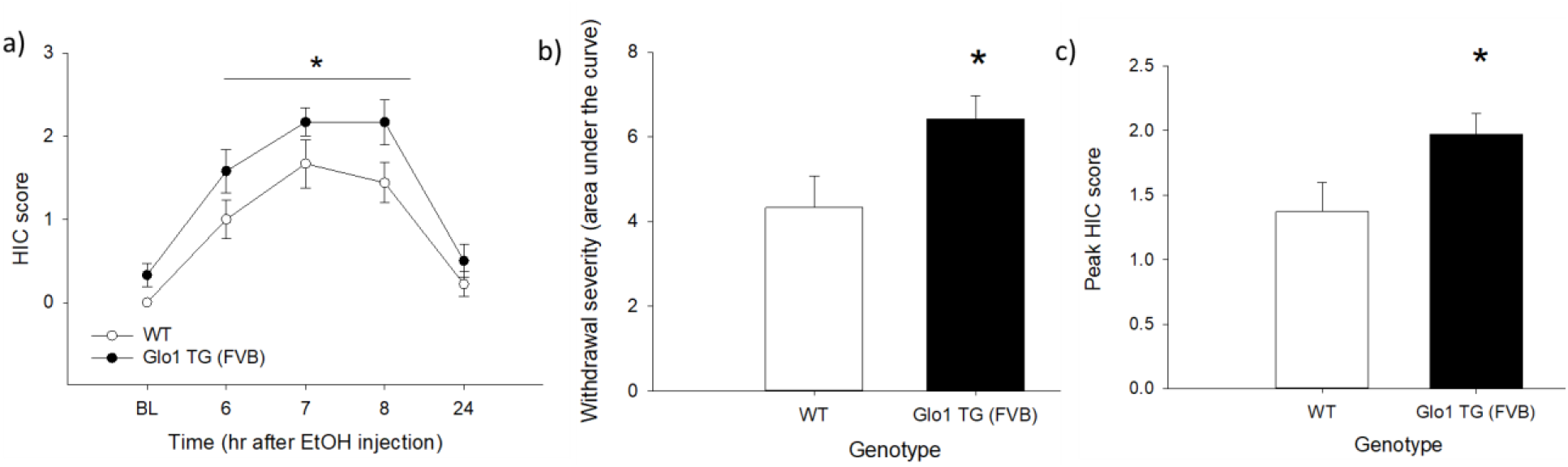
Acute ethanol withdrawal as assessed by handling-induced convulsions (HICs) in wild type (WT) and *Glo1* overexpressing (TG) mice on an FVB genetic background. Panel **a** shows time course of withdrawal, with baseline HIC measurements made immediately before ethanol injection. There was a main effect of genotype, with *Glo1* overexpressing mice having higher HIC scores across the time course that WT mice. *Glo1* overexpressing mice also showed greater cumulative seizure susceptibility (area under the curve [AUC], panel **b**), and higher peak HIC scores than WT littermates (panel **c**). * indicates p<0.05 N=9-12/genotype.

### Experiment 3: GLO1 inhibitor effects on chronic ethanol withdrawal

Figure 3 shows withdrawal HIC measures after one or two cycles of chronic ethanol treatment. No animals in the non-withdrawal control groups (saline-vehicle and saline-pBBG) showed any seizure activity (0 scores for all animals at all time points), so only the data from the withdrawal group are reported here. Analysis with mixed-model ANOVA of HIC time course found a main effect of time following a single ethanol cycle (**Fig 3a**: *F*_3.19_,70.06=3.24, p=0.025) and a trend toward a main effect of genotype (*F*_1,22_=3.28, p=0.084), but it did not reach the level of statistical significance. There was also no significant treatment x time interaction (p>0.1). After two cycles of ethanol exposure, there were main effects of both time and treatment (**Fig 3b**: *F*_2.77,61.02_=4.44, p=0.008 and *F*_1,22_=5.38, p=0.03, respectively). The time x treatment interaction was not statistically significant (*F*_2.77,61.02_=2.28, p=0.093).

**Figure 3.**
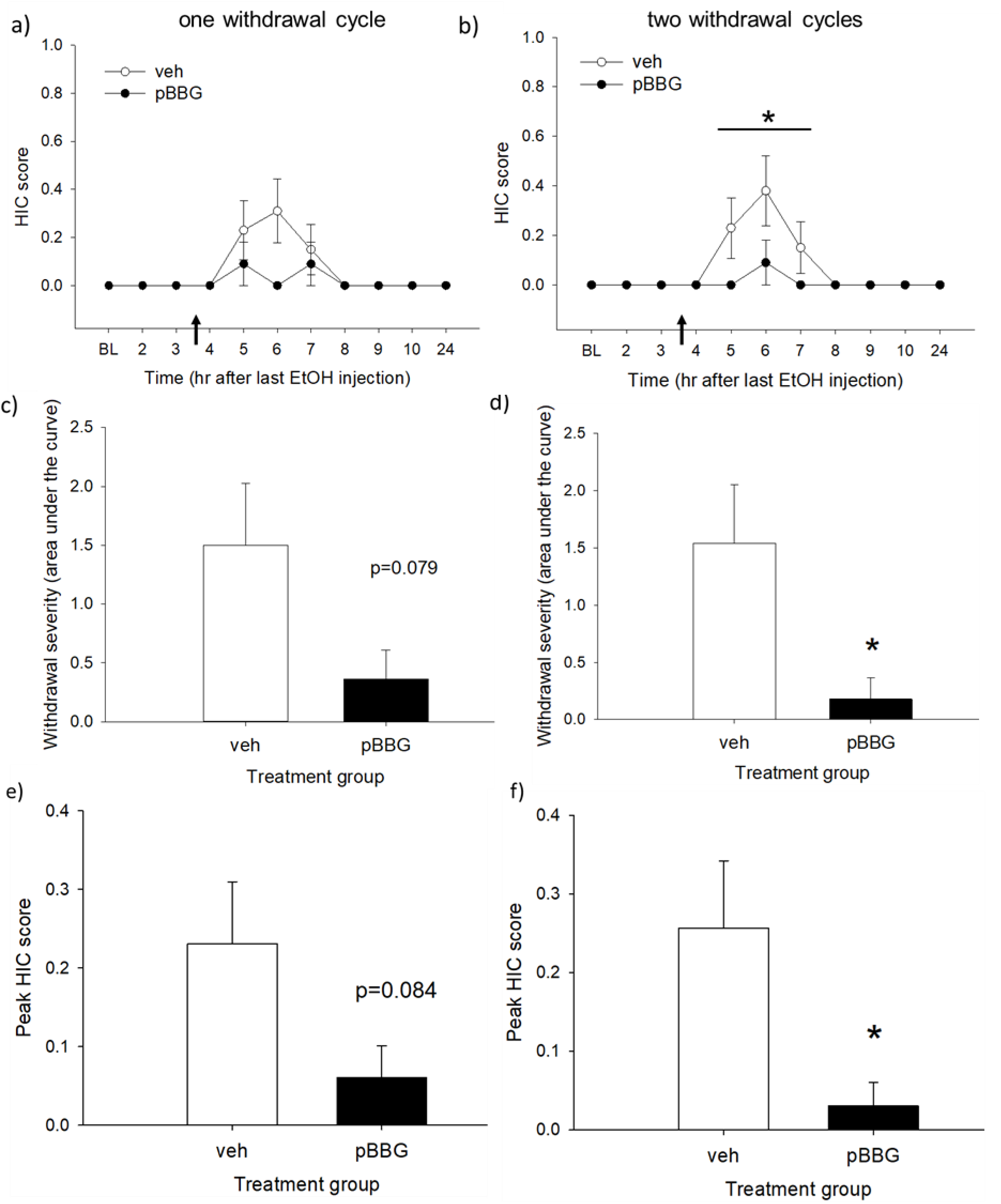
HIC time courses (a & b), area under the HIC curve (c & d) and peak HIC scores (e & f) after one (left column) and two (right column) withdrawal cycles. Baseline HIC measurements were made before the start of each 10-day ethanol exposure period. Injections of GLO1 inhibitor (pBBG, 25 mg/kg) or vehicle were given at 3.5 hr after the final ethanol injection (indicated by arrows). * indicates statistically significant difference from vehicle-treated mice (p<0.05). N=11-13/treatment group.

AUC was calculated for hours 4-10, as this was the time range when the GLO1 inhibitor treatment was expected to have an effect. Analysis of area under the HIC curve showed that following one cycle of ethanol, there was a trend toward lower HIC AUC in the inhibitor-treated group compared to vehicle (**Fig 3c**: *F*_1,22_=3.38, p=0.079), but it was not statistically significant. There was also not a significant main effect of treatment group on peak HIC score (Fig 3d; *F*_1,22_=3.28, p=0.084) or latency to peak HIC (*F*_1,6_,=0.19, p= 0.68, data not shown) after one cycle of ethanol. In contrast, after two cycles of ethanol exposure, there were significant effects of treatment group on both HIC AUC (**Fig 3e**) and peak HIC score (**Fig 3f**) with GLO1 inhibition significantly reducing both measures (*F*_1,22_=5.38, p=0.03 for both). Latency to peak HIC again did not differ between the groups (*F*_1,6_=0.56, p=0.482, data not shown).

Because of the low overall withdrawal scores in the vehicle-treated animals (HIC scores ranged from 0-1) and the potential of a floor effect, one-sample t-tests were used to determine whether HIC AUC in each cycle was significantly different from 0 (AUC for non-withdrawal control groups). HIC AUC was significantly different from 0 following both one and two cycles of ethanol treatment (cycle 1: *t*(12)=2.84, p=0.015; cycle 2: *t*(12)=2.99, p=0.011; Bonferroni corrected α=0.025), indicating that there was measurable withdrawal in the vehicle-treated group which was reduced by pBBG treatment.

## 4. Discussion

In these experiments, we showed that overexpression of the gene *Glo1* can produce increased ethanol withdrawal severity as assessed by HICs. Furthermore, treatment of wild type animals with a GLO1 inhibitor had the opposite effect, namely attenuating ethanol withdrawal seizure severity in mice undergoing a chronic ethanol exposure procedure. Taken together, these results provide evidence that GLO1 activity modulates CNS hyperexcitability during ethanol withdrawal in mice.

Experiment 1 showed a weak trend towards increased HIC scores in mice overexpressing *Glo1* on a B6 background, but this difference did not reach the threshold for statistical significance. In Experiment 2, however, overexpression of *Glo1* on an FVB background was found to significantly increase both HIC AUC and peak HIC score. B6 mice have been shown previously to be largely resistant to ethanol withdrawal seizures [20, 38–40] and this is consistent with our findings here where most animals had low or no seizure activity and the observed HIC scores only ranged from 0-2 (Table S1). In contrast, mice in Experiment 2 showed much greater seizure activity, with HIC scores ranging from 0-4. Therefore, it is possible that genetic background is an important mediator of *Glo1s* effects on ethanol withdrawal. The two transgenic lines also differ in their number of copies of *Glo1* (the B6 line has 17 copies, whereas the FVB line has 48 copies), and this could also contribute to the differences in withdrawal potentiation between the lines.

GLO1 activity has not been previously examined as a potential mediator of ethanol withdrawal severity. Chronic ethanol produces significant neuroadaptations, particularly in the GABA and glutamate systems, which underlie some of the symptoms of ethanol withdrawal (e.g. [30, 41–43]). Previous work in the lab has demonstrated that the *Glo1* transgenic mice have increased GLO1 enzymatic activity and decreased brain MG levels [23]. We have also shown that MG can act directly at GABA_A_ receptors [23]. One possible explanation for the effects of GLO1 manipulations on ethanol withdrawal seizures, therefore, may be subsequent downstream changes in MG levels and activity at GABA_A_ receptors. We have shown previously that MG treatment reduces both the severity and duration of pharmacologically induced seizures, as well as EEG measures of seizure activity [29]. Additionally, both acute and chronic ethanol treatment have been shown to decrease brain glucose metabolism in humans and rodents [44–46], and brain glucose metabolism increases during abstinence [47]. It is possible that these metabolic alterations lead to changes in MG brain levels during withdrawal which might be reversed by GLO1 inhibition. However, it should be noted that we did not directly measure MG levels during withdrawal in these studies, and therefore these possibilities are speculative. Direct measurement of MG brain levels during withdrawal will be an important step for determining how the glyoxalase system specifically may be altered by ethanol treatment and subsequent withdrawal. GLO1 effects on negative affective changes during protracted abstinence should also be examined, since many of these changes (e.g. increased anxiety- and depression-like behavior) have been seen with genetic and pharmacological manipulation of GLO1 activity in nondependent mice [23–25].

The finding in Experiment 3 that pBBG treatment only produced a trend toward an effect following one cycle of ethanol exposure but led to significant decreases in HIC severity after two cycles, suggests that GLO1 inhibition may be more effective in animals with a greater dependence history. This is consistent with previous findings where pBBG was more effective at reducing ethanol drinking in dependent rats compared to non-dependent rats [28]. In the present study there was not a significant ethanol cycle x treatment group interaction for either AUC or peak HIC score (data not shown), so we cannot state conclusively what may be driving the increased efficacy seen with two ethanol withdrawal cycles. There is considerable evidence that ethanol withdrawal severity increases with repeated withdrawal experience (“kindling” effect) (e.g. [48–50]). It could be that withdrawal severity increased modestly with the repeated ethanol cycles, thereby eliminating any floor effect and making it easier to see a statistically significant reduction in seizure activity following pBBG treatment. Alternatively, it is possible that pBBG is more effective in animals with a greater ethanol history due to the subsequent neuroadaptations, or a combination of these factors.

There are some limitations of the present studies that should be noted. First, potential sex differences were only directly examined in Experiment 1, and the number of animals per sex per genotype likely provided limited power for identifying interactions with sex. However, we did see significant and consistent effects of GLO1 manipulations in both male and female animals in Experiments 2 and 3, even though the sexes were not directly compared in these experiments. Mice in Experiment 2 were also older on average than mice in the other two experiments, and potential age effects cannot be ruled out with the current experimental design. However, the similar direction of effects of *Glo1* overexpression seen in Experiments 1 and 2 using transgenic mice on two different genetic backgrounds, the parallel effect of a GLO1 inhibitor in wild type animals, and the consistency in findings in both acute and chronic withdrawal models suggest that the role of GLO1 in ethanol withdrawal severity generalizes across multiple populations and conditions. Additionally, although GLO1 inhibition reduced HICs, it did not return animals completely to baseline levels. Testing a wider range of pBBG doses could help to determine whether there is an optimal dose that can fully eliminate withdrawal seizures. Finally, the overall level of withdrawal severity seen in Experiments 1 and 3 was low, likely due to the use of B6 mice, and future studies in this genotype might benefit from a more extensive ethanol exposure method to induce greater dependence (e.g. chronic-intermittent ethanol vapor exposure).

In conclusion, we provided converging genetic and pharmacological evidence that GLO1 activity mediates ethanol withdrawal seizure susceptibility. In mice, overexpression of *Glo1* contributes to increased withdrawal seizure severity, and inhibition of GLO1 attenuates withdrawal seizure severity. GLO1 inhibition could therefore potentially be a useful strategy for treatment of neural hyperexcitability during acute withdrawal. Further studies will be needed to determine how the glyoxalase system is altered by chronic ethanol exposure, and whether GLO1 inhibitors may have utility for treating negative affective changes seen during protracted abstinence.

## Author Contributions

Conceptualization, AMB-L, AL, and AAP; Formal analysis, AMB-L and AL; Funding acquisition, AAP; Investigation, AMB-L and AL; Project administration, AMB-L; Supervision, AAP; Writing – original draft, AMB-L; Writing – review & editing, AMB-L, AL, and AAP. All authors approved the final version of the manuscript for submission.

## Funding

This work was supported by NIH-NIAAA grant R01 AA026281. AMB-L was supported by NIH-NIAAA grants F32 AA025515 and K99 AA07835.

## Conflicts of Interest

The authors declare no conflict of interest.

---

**Supplemental Table S1:** Number of animals per maximum HIC score for each experiment

|  | Group | No HICs (0) | 1 | 2 | 3 | 4 |
| --- | --- | --- | --- | --- | --- | --- |
| **Experiment 1** | WT | 5 | 3 | 0 | 0 | 0 |
|  | TG (B6) | 3 | 4 | 1 | 0 | 0 |
| **Experiment 2** | WT | 1 | 1 | 6 | 1 | 0 |
|  | TG (FVB) | 0 | 0 | 7 | 3 | 2 |
| **Experiment 3 (1 cycle)** | Vehicle | 7 | 6 | 0 | 0 | 0 |
|  | pBBG | 9 | 2 | 0 | 0 | 0 |
| **Experiment 3 (2 cycles)** | Vehicle | 6 | 7 | 0 | 0 | 0 |
|  | pBBG | 10 | 1 | 0 | 0 | 0 |

**Figure S1.**
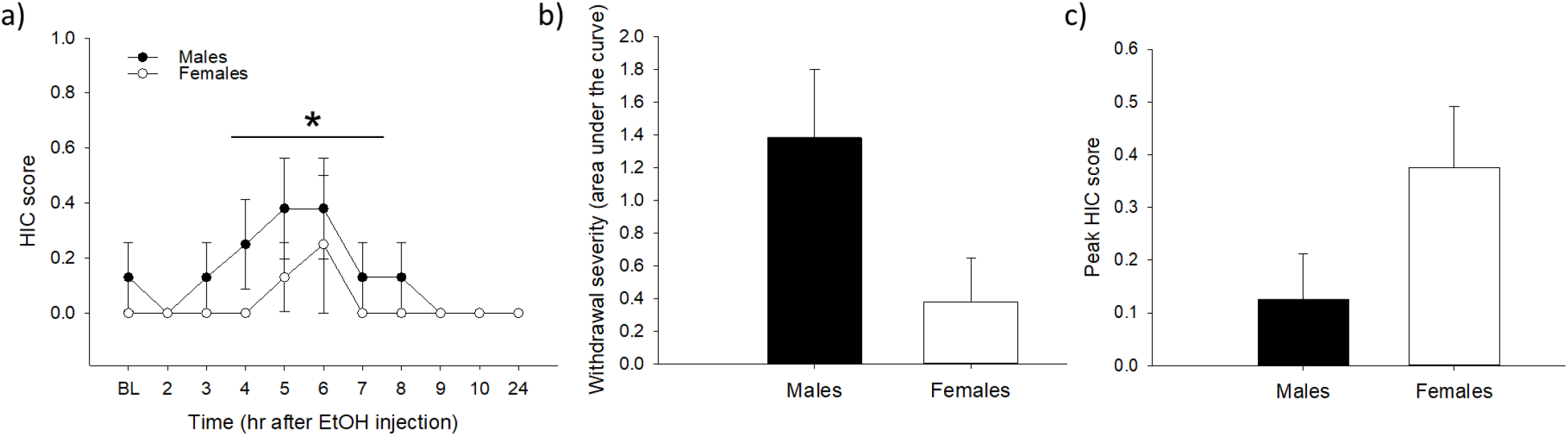
Acute ethanol withdrawal measures for male and female mice from Experiment 1 collapsed on genotype. Panel **a** shows time course of withdrawal, with baseline HIC measurements made immediately before ethanol injection. There was a main effect of sex, with male mice having higher HIC scores across the time course than female mice. Panel **b** shows cumulative HIC scores (area under the curve, AUC) for each sex. There was a non-significant increase in HIC AUC in males compared to females (p=0.064). Panel **c** shows peak HIC score for each sex. N=8/genotype. * p<0.05

## Notes

### Competing Interest Statement

The authors have declared no competing interest.

